# Cell line-dependent effects of spheroid formation method on drug response in melanoma models

**DOI:** 10.64898/2026.05.12.724514

**Authors:** Akvilė Žilytė, Vilma Petrikaitė

## Abstract

In this study, we evaluated the impact of different in vitro 3D culture modelling methods on the activity of doxorubicin (DOX) and 5-fluorouracil (5-FU) in human melanoma spheroids. Human melanoma A375 and IGR39 spheroids were generated using the hanging drop and non-adhesive surface methods. Spheroid growth dynamics were assessed by measuring changes in spheroid diameter. To compare the effects of anticancer drugs in spheroids of different sizes, spheroids of approximately 200 and 400 µm were formed. Drug activity was evaluated based on spheroid growth and cell viability using the MTT assay. A375 spheroids formed using the non-adhesive surface method were more sensitive to DOX than spheroids formed using the hanging drop method. In smaller A375 spheroids, 10 µM 5-FU reduced cell viability more effectively in spheroids formed using the hanging drop method. In contrast, IGR39 spheroids formed by the hanging drop method were more resistant than those formed on a non-adhesive surface. However, in IGR39 spheroids, the effects of DOX and 5-FU on growth and viability did not significantly differ between formation methods. In conclusion, A375 spheroid growth was not significantly influenced by the formation method, whereas IGR39 spheroid growth depended on the method used. A375 spheroids formed on non-adhesive surfaces were more sensitive to DOX, whereas 5-FU activity depended on drug concentration and spheroid size. In IGR39 spheroids, the effects of DOX and 5-FU on growth and viability were largely independent of the spheroid formation method. Based on these results, it can be concluded that the researchers should carefully select the spheroid formation method for their studies, as this may influence the results of the tested compound’s effect on their size and viability.

## 1. Introduction

Recently, studies using three-dimensional (3D) cell culture models have gained increasing attention compared to traditional two-dimensional (2D) cultures, as they better mimic the tumour microenvironment and in vivo conditions [1–3]. Before initiating clinical studies, preclinical studies are conducted to evaluate the effects of the developed pharmaceutical substances using in vitro models and animal experiments. However, animal models are expensive and do not always yield successful results [4]. In addition, the 3R principle (replacement, reduction, refinement) promotes reducing animal use in research and improving animal welfare. As a result, in vitro models are increasingly applied in research [5].

In vitro models are generally divided into 2D and 3D cell culture models. Conventional 2D culture models are unable to reproduce the heterogeneous properties characteristic of the tumour microenvironment [2]. In contrast, 3D cell culture models more closely resemble real tissues and are therefore used to create more accurate and complex tumour models [1]. Cells in 2D cultures are often more sensitive to drugs than cells in 3D cultures because they grow in a flat plane, lose their natural spatial structure and may gradually die. In contrast, cells in 3D cultures maintain their interconnections, spatial structure, and viability [3].

Among 3D cultures, spheroids are one of the simplest and most widely used models. Spheroids contain multiple zones with different conditions, replicating in vivo tumour characteristics such as cell hypoxia, nutrient gradients, growth kinetics, and gene expression [6]. Spheroids are usually round, with surface points equidistant from the centre, allowing to understand the concentration gradients of oxygen, nutrients, and metabolites in tumours. Based on cell composition, spheroids are classified as homotypic (composed of tumour cells) or heterotypic spheroids (containing fibroblasts, immune cells and cancer cells) [6]. Due to these properties, spheroids are valuable tools for studying cancer biology and anticancer drug activity [2,7].

Various spheroid formation techniques have been developed [8]. However, comparing results between studies remains challenging because different methods often generate spheroids with distinct morphological and physiological properties [2,3]. One of the most important characteristics is spheroid size. Spheroids larger than 400 μm develop a hypoxic core that resembles conditions found in tumours [9,10]. Hypoxia activates signalling pathways that maintain cell viability and promote stable expression of hypoxia-inducible factors (HIFs), which stimulate angiogenesis, glycolysis, and pH regulation changes. Consequently, spheroids of different sizes exhibit differences in hypoxia, nutrient gradients, and drug penetration, which may influence experimental outcomes and complicate comparisons between studies [2]. Nevertheless, insufficient attention is often paid to the spheroid formation method itself when interpreting or comparing published data.

Therefore, in this study, we aimed to evaluate how different 3D spheroid formation methods influence melanoma spheroid growth and sensitivity to anticancer drugs. Human melanoma was selected as the experimental model because melanoma is a highly aggressive skin cancer with a strong tendency to metastasize to the lymphatic system [11]. According to the European Cancer Information System (ECIS), melanoma affected approximately 30 out of 100,000 people in Europe in 2022 [12]. Furthermore, melanoma Incidence and mortality rates continue to increase worldwide, highlighting the need for improved experimental models and more effective therapeutic strategies [12].

Two commonly used spheroid formation models were selected: the hanging drop method and the non-adhesive surface method [13]. The hanging drop method is one of the oldest and most frequently used techniques due to its simplicity and the possibility of selective monitoring [3]. Through the combined effects of surface tension and gravitational force, cells aggregate and form spheroids. This method enables the generation of uniformly sized spheroids and allows precise control of environmental conditions [14]. The non-adhesive surface method is also often used because of its simplicity. Non-adhesive surfaces prevent cell adhesion, thereby promoting cell-cell interactions and facilitating spheroid formation. In addition, spheroid size can be regulated by adjusting the initial cell seeding density [15].

Two melanoma cell lines, IGR39 and A375, were used in this study. Both cell lines carry the BRAFV600E mutation, which promotes cell proliferation and increases resistance to apoptosis [16]. However, these cell lines differ in their expression of estrogen receptor β (ERβ). Estrogen receptors are divided into ERα, associated with proliferative activity, and ERβ, which is linked to anticancer effects. Therefore, the influence of estrogen receptors on tumour behaviour may depend on their relative expression in a specific tumour site in the body. ERβ is virtually undetectable in IGR39 cells, whereas A375 cells express this receptor, suggesting potential differences in sensitivity to anticancer drugs [17].

The study focused on the effects of doxorubicin (DOX) and 5-fluorouracil (FU), which exhibit different mechanisms of anticancer activity. DOX is a widely used cytotoxic chemotherapeutic agent that induces cell death mainly through DNA intercalation and topoisomerase II inhibition [18–20]. In contrast, 5-FU is a cytostatic drug that interferes with DNA and RNA synthesis by inhibiting thymidylate synthase [21,22]. Due to their distinct mechanisms of action, both drugs were selected to evaluate how different spheroid formation methods influence melanoma cell responses to anticancer treatment.

## 2. Materials and Methods

### 2.1. Cell culture cultivation

IGR39 (Deutsche Sammlung von Mikroorganismen und Zellkulturen GmbH, DSMZ) and A375 (the American Type Culture Collection, ATCC) cell lines were cultured at 37°C in a 5% CO_2_ atmosphere in Dulbecco’s Modified Eagle GlutaMAX cell culture medium (DMEM GlutaMAX, Gibco) supplemented with 10% fetal bovine serum (FBS, Gibco) and 1% antibiotic solution (10,000 IU/ml penicillin and 10 mg/mg streptomycin, Gibco), until 70% confluence. The cell medium was then aspirated, and the cells were washed with phosphate-buffered saline (PBS, pH 7.4, Gibco). After aspirating the PBS solution, cells were trypsinised using TrypLE Express (Gibco). Detached cells were resuspended in cell culture medium and centrifuged at 1000 rpm for 4 min. After discarding the supernatant, the cells were resuspended in fresh medium. Cell lines were cultured until passage 20.

### 2.2. Spheroid formation using the hanging drop method

To form spheroids by the hanging drop technique, a cell suspension was prepared as described in section 2.1. Cells were counted using a Neubauer chamber (Sigma-Aldrich). Then, serial dilutions of cell suspension were prepared (1 × 10^5^ cells/ml, 5 × 10^4^ cells/ml, 2.5 × 10^4^ cells/ml, and 1.25 × 10^4^ cells/ml) in a medium containing 0.024% methylcellulose (MC, 4000 cP, Sigma-Aldrich). The drops of prepared cell suspension were placed on the inverted plate lid. Then, the lid was quickly flipped back to its normal position and placed on the bottom lid of the plate with 3 ml of sterile water. The plates were incubated until spheroids were formed, and throughout the whole experiment at 37°C in a 5% CO_2_ atmosphere.

### 2.3. Formation of spheroids on a non-adhesive surface

The 96-well plate was coated with a 2% agarose (molecular biology grade, Roth). Then, 100 µl of the cell suspension at serial dilutions (2 × 10^4^ cells/ml, 1 × 10^4^ cells/ml, 5 × 10^3^ cells/ml, 2.5 × 10^3^ cells/ml) was added into each well. The plates were incubated until spheroids were formed, and throughout the whole experiment at 37°C in a 5% CO_2_ atmosphere.

### 2.4. Investigation of spheroid growth

Photos of the spheroids formed were captured at 10× magnification using a phase-contrast regimen by an inverted microscope Olympus IX73 (Olympus Europe Holding GmbH). Eight spheroids were used in each group (eight technical repetitions in one biological experiment). The spheroids were photographed every other day for 10 days. The medium was changed every 4 days. The images obtained were analysed using ImageJ 1.53t (National Institutes of Health, Bethesda, MD, USA). The diameter of the spheroid was calculated from the spheroid area. The analysis was performed using Microsoft Office Excel 2024 (Microsoft Corporation) software.

### 2.5. Determination of the effect of doxorubicin and 5-fluorouracil on spheroid growth and viability

The effects of DOX (98%, Abcam) and 5-FU (≥ 98%, Abcam) were studied on spheroids formed using previously described techniques. For these experiments, spheroids of two different diameters were selected: 200 µm and 400 µm, in order to compare the activity of anticancer drugs depending on the hypoxia present in them. Before adding the tested drugs, spheroids formed by the hanging drop technique were transferred to an agarose-coated plate. Then, 100 µl of 2 and 10 µM DOX and 2 and 10 µM 5-FU solutions were added to the wells with spheroids. A medium containing 0.2% DMSO (≥ 99%, Chemlab) was added to the control wells. Eight spheroids were used in each group (eight technical repetitions in one biological experiment). The spheroids were incubated at 37 °C in a 5% CO2 environment. The medium was changed every 4 days, and the photos of spheroids were captured every other day for 8 days.

After 8 days of incubation with anticancer drugs, viability was assessed using the 3-(4,5-dimethylthiazol-2-yl)-2,5-diphenyltetrazolium bromide (MTT) assay. On the last day of the experiment, the medium with drugs was carefully aspirated and replaced with a fresh medium containing 0.5 mg/ml MTT (97%, Sigma-Aldrich). After 12-14 hours, the spheroids were transferred to a new 96-well plate and washed with PBS. Then, the formazan crystals were dissolved in DMSO. The absorbance of the solution was measured by a Multiskan GO Microplate spectrophotometer (Thermo Fisher Scientific) at wavelengths of 570 nm and 630 nm. The analysis was performed using Microsoft Office Excel 2024 software.

### 2.6. Statistical data evaluation

The research data were analysed using Microsoft Office Excel 2024 software, evaluating averages and relative standard deviations from three independent biological experiments. The averages were compared using Student’s t-test. Differences were considered statistically significant when the significance level was p < 0.05.

## 3. Results and discussion

### 3.1. Comparison of the growth dynamics of human melanoma spheroids formed using different methods

A375 cells by both spheroid formation techniques on the second day, formed compact and round structures (Figures 1A and B). At the beginning of the experiment, the spheroids formed using the hanging drop technique were 1.2-1.3 times smaller than those formed using the non-adhesive surface technique (p < 0.05) (Figures 1C, 1D, 1E, and 1F).

**Figure 1.**
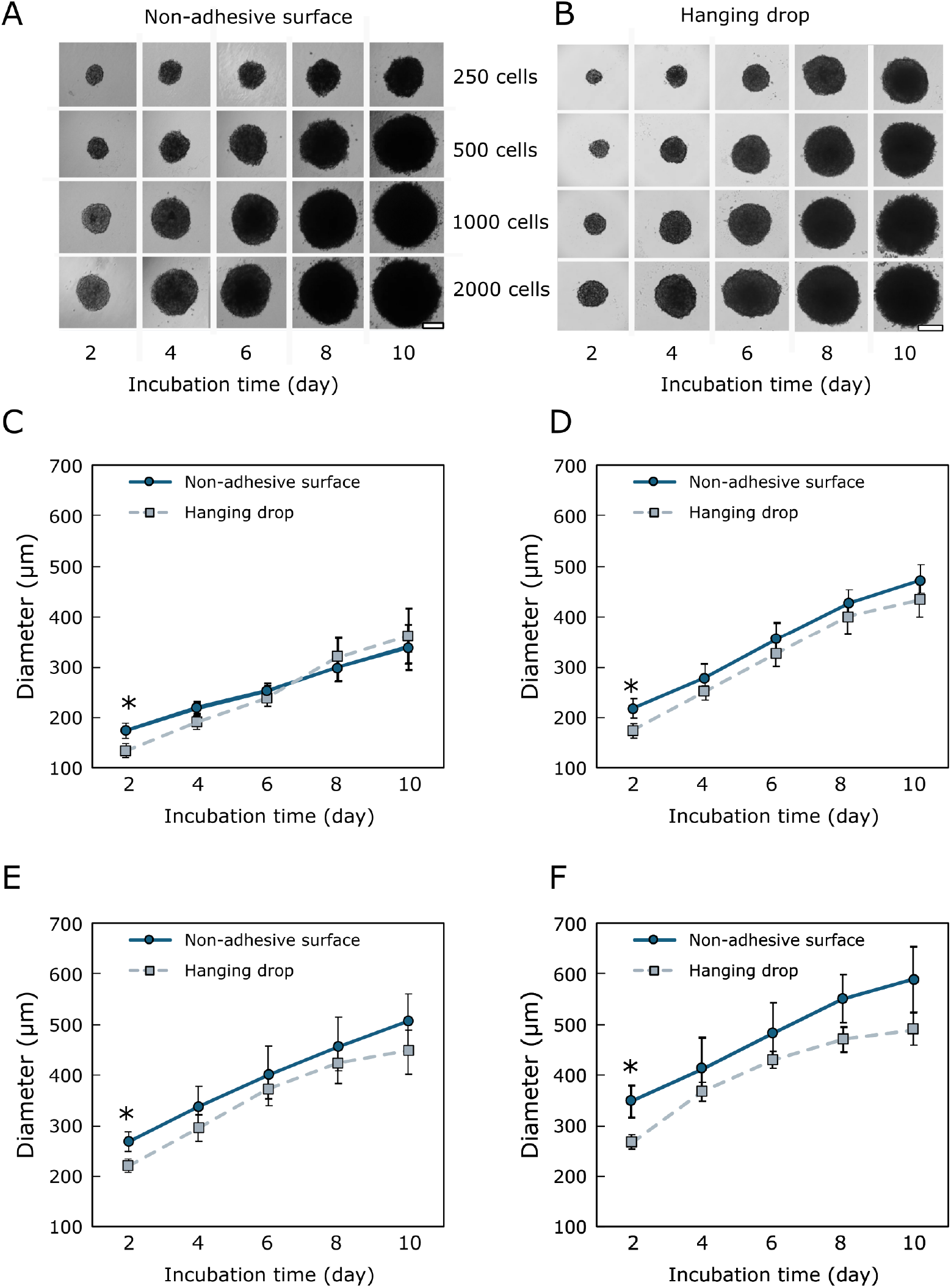
Growth of A375 spheroids for 10 days. Photos of A375 spheroids formed using the non-adhesive surface (**A**) and hanging drop (**B**) techniques. Diameter of A375 spheroids formed from 250 (**C**), 500 (**D**), 1000 (**E**), and 2000 (**F**) cells. Scale bar 200 µm, *p < 0.05, statistically significant when comparing spheroids formed by different methods.

The diameter of A375 spheroids formed by the hanging drop technique ranged from 137.0 ± 6.4 µm (on the second day of incubation) to 486.4 ± 31.7 µm (on the tenth day of incubation). On the second day of incubation, the diameter of the spheroids was approximately 140 µm when formed from 250 cells; 170 µm from 500 cells; 220 µm from 1000 cells, and 260 µm from 2000 cells. Spheroids formed from 250 and 500 cells increased by 2.5-fold after ten days of incubation (Figures 1C and 1D). Spheroids formed from 1000 and 2000 cells grew more slowly and a 2-fold increase in diameter after ten days of incubation (Figures 1E and 1F).

The size of A375 spheroids formed using the non-adherent plate method varied from 175.4 ± 11.7 µm to 585.3 ± 65.3 µm throughout the experimental period. After two days of incubation, compact spheroids developed a diameter of approximately 180 µm when formed from 250 cells, 220 µm from 500 cells, 270 µm from 1000 cells, and 340 µm from 2000 cells. Spheroids formed from 250 and 500 cells increased approximately 2-fold after ten days of incubation (Figures 1C and 1D). Spheroids formed from 1000 and 2000 cells grew more slowly and increased by 1.8-fold after ten days of incubation (Figures 1E and 1F).

The results indicate that the A375 spheroids formed using both methods grew at a similar rate, although the spheroids formed on a non-adhesive surface were larger at the beginning. No significant differences were observed at later time points. Smaller spheroids (250 and 500 cells) grew 2-2.5-fold faster, while larger spheroids (1000 and 2000 cells) grew 1.8-2-fold faster than spheroids formed by hanging drop.

For further studies of the activity and transport of anticancer drugs, different cell concentrations were selected. After an appropriate incubation time, 200 µm and 400 µm spheroids with different oxygen levels were produced.

IGR39 spheroids generated using the hanging drop and non-adhesive surface methods formed at different time points (Figure 2). On a non-adhesive surface, the spheroids formed on the second day (Figure 2A), while in the hanging drops, spheroidal structures developed on the fourth day (Figure 2B). On the second day of incubation, the IGR39 spheroids generated from 1000 and 2000 cells using the non-adhesive surface method were round, compact, and retained this shape throughout the study. Meanwhile, the spheroids developed from a smaller number of cells on a non-adhesive surface method became round and compact only on the sixth day of incubation (Figure 2A), while those formed using the hanging drop method remained irregular in shape throughout the experiment (Figure 2B).

**Figure 2.**
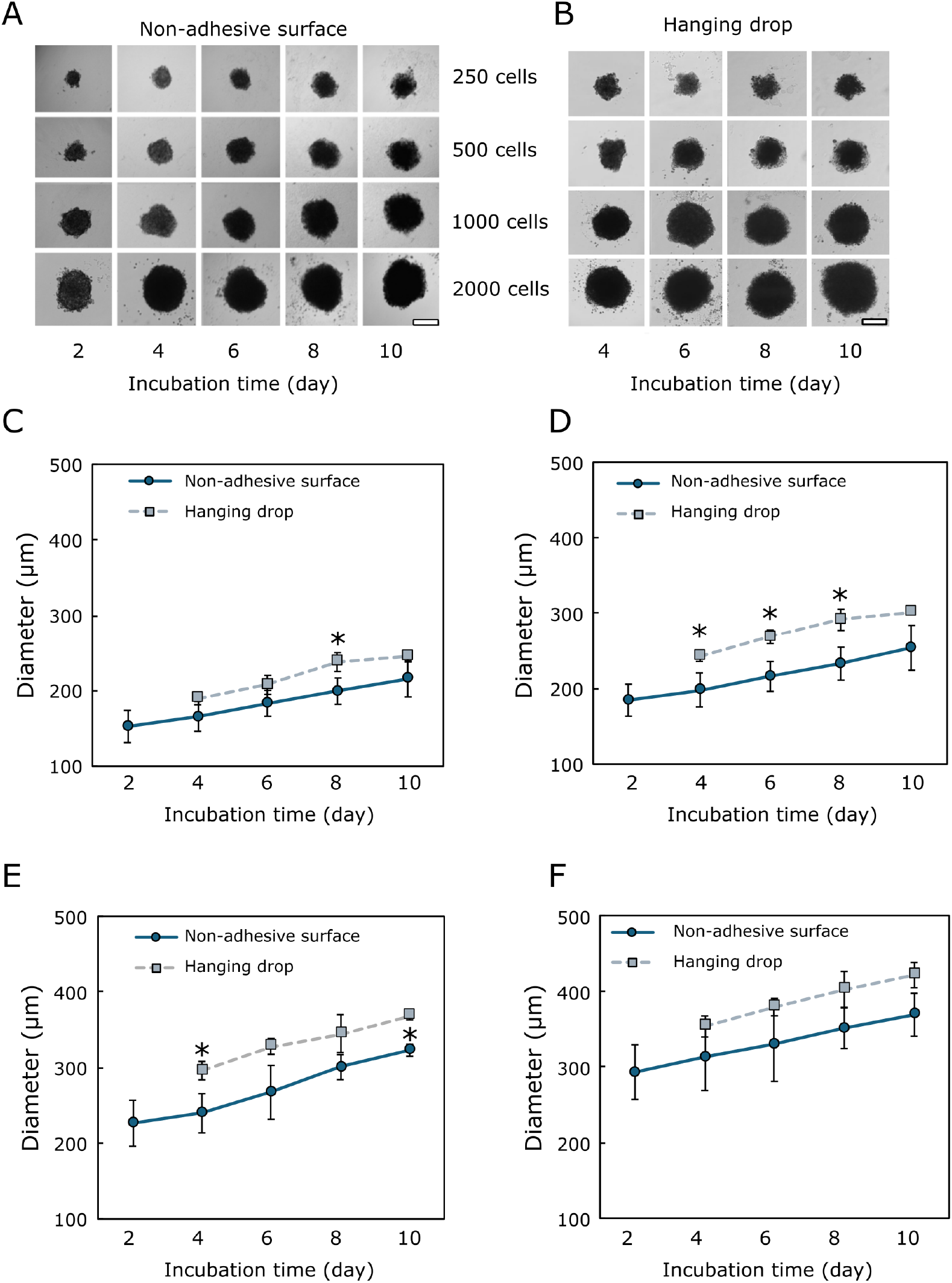
Growth of IGR39 spheroids for 10 days. Photos of IGR39 spheroids formed using the non-adhesive surface (**A**) and hanging drop (**B**) techniques. Diameter of IGR39 spheroids formed from 250 (**C**), 500 (**D**), 1000 (**E)**, and 2000 (**F**) cells. Scale bar 200 µm, *p < 0.05, statistically significant when comparing spheroids formed by different methods.

The diameter of IGR39 spheroids generated by the hanging drop method ranged from 189.3 ± 7.8 µm to 421.3 ± 16.7 µm throughout the experiment (Figures 2C-F). After four days of incubation, compact spheroids developed with a diameter of approximately 190 µm when formed from 250 cells; 220 µm from 500 cells; 300 µm from 1000 cells, and 350 µm from 2000 cells. Spheroids formed by the hanging drop method from 250, 500, 1000, and 2000 cells showed an approximately 1.2-fold increase over ten days.

The size of IGR39 spheroids formed on a non-adhesive surface ranged from 152.6 ± 21.3 µm to 368.7 ± 28.4 µm (Figures 2C-F). On the second day, the spheroid diameter was approximately 150 µm when formed from 250 cells; 185 µm from 500 cells; 230 µm from 1000 cells, and 300 µm from 2000 cells.

Throughout the experiment duration, the diameter of IGR39 spheroids formed from 500 and 1000 cells on a non-adhesive surface was approximately 1.1-fold smaller than that of spheroids generated by the hanging drop method (Figures 2D-F). However, statistically significant differences between methods were observed in a group of spheroids formed from 500 cells after four, six, and eight days of incubation, and in a group of spheroids formed from 1000 cells after four and ten days. In contrast, no significant size differences were observed in spheroids generated from 250 or 2000 cells. For further studies of anticancer drug activity and viability, two cell concentrations were selected to obtain spheroids of the desired size: 250 and 2000 cells for the hanging drop technique, and 500 and 2000 cells using a formation on a non-adhesive surface.

Comparing the formation and growth dynamics of A375 and IGR39 spheroids, it was revealed that both cell lines formed spheroids on a non-adhesive surface by day two. IGR39 spheroids using the hanging drop technique were developed later (after four days) (Figure 1 and Figure 2). By day ten, A375 spheroids formed in hanging drops were approximately 1.4-fold larger than IGR39 spheroids when generated from 250 and 500 cells and about 1.1-fold larger when generated from 1000 and 2000 cells.

Raghavan et al. formed breast and ovarian cancer spheroids using the hanging drop and non-adherent plate methods. Unlike in our study, the spheroids of both lines were formed from a smaller number of cells and grown for seven days. However, similarly to our study, at the end of the experiment, the spheroids of both lines formed using the hanging drop method were about 3 times smaller than the spheroids formed using the non-adherent plate method, regardless of the cell seeding density [1].

Jeong et al. formed colon cancer spheroids by the hanging drop technique and also on a non-adhesive surface. By using both methods, spheroids were formed from 300 to 20,000 cells. Spheroids formed using the hanging drop technique were grown for 14 days, while those formed on a plate were grown for 35 days. On the 14th day, the spheroids formed on a non-adhesive surface were twice as large as those formed using the hanging drop method. The size of the spheroids was determined by the formation method. It is believed that by improving the hanging drop method, it is possible to grow spheroids with a larger diameter and use them for a long time [23].

Generally, the growth dynamics of human melanoma A375 spheroids in our study did not depend on the formation technique. The size of the spheroids differed only at the beginning of the study, but in later periods, the size of the spheroids was similar. Meanwhile, the size of the IGR39 line spheroids depended on the initial number of cells used. When formed from 250 and 2000 cells, there was no difference in the size of the spheroids after they had formed and during their growth until the tenth day. When IGR39 spheroids were formed from 500 and 1000 cells, the spheroids formed using the hanging drop method remained larger than those formed on a non-adhesive surface over a period of ten days.

### 3.2 The influence of spheroid formation technique on their sensitivity to DOX and 5-FU

Generally, the sensitivity of human melanoma spheroids exposed to cytotoxic (DOX) and cytostatic (5-FU) compounds varied depending on the formation technique chosen, the initial number of cells, and the concentration of the tested compound.

A375 spheroids, regardless of the formation method and spheroid size, showed a reduction in diameter after eight days of incubation with 2 µM and 10 µM DOX compared to the control (Figure 3). Smaller (200 µm diameter) A375 spheroids formed by the hanging drop technique were approximately 1.3-fold smaller after eight days of incubation with 10 µM DOX compared to the control. In contrast, spheroids produced in the non-adhesive surface exhibited an approximately 2.3-fold reduction compared to the control (Figure 3A). Similarly, smaller spheroids generated using the non-adhesive surface method and incubated with 2 µM DOX had a diameter that was 2.3-fold smaller. Meanwhile, spheroids formed using the hanging drop method did not show a statistically significant change compared to the control (Figure 3A).

**Figure 3.**
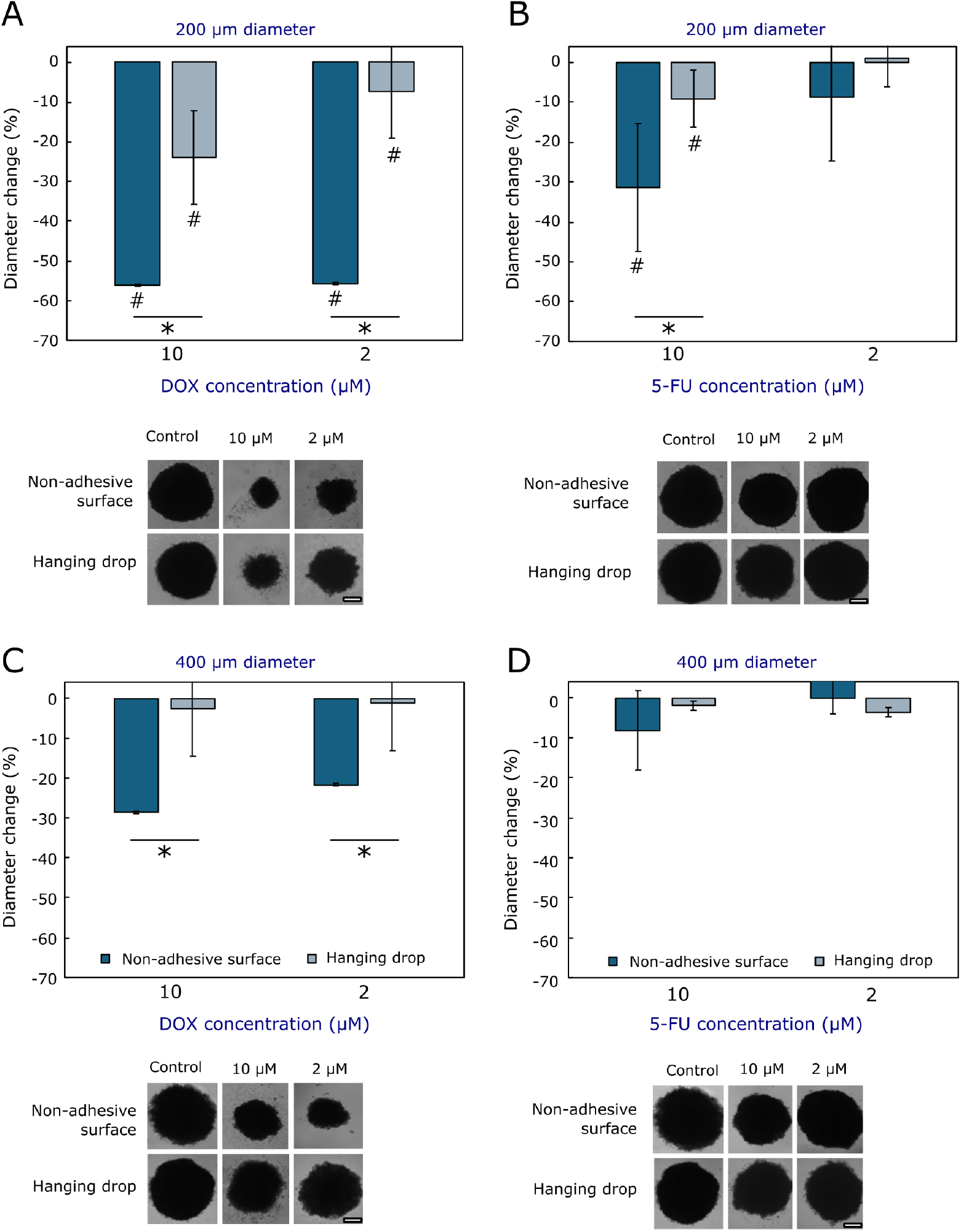
Change in diameter of 200 µm (**A, B**) and 400 µm (**C, D**) A375 spheroids after 8 days of incubation with DOX and 5-FU; *p < 0.05, statistically significant when comparing spheroids formed by different methods, #p < 0.05, statistically significant when compared to the control. Representative photos of A375 spheroids after 8 days of incubation with tested compounds are provided below each graph; scale bar corresponds to 200 µm.

A similar trend was observed when smaller spheroids were incubated with 5-FU. Spheroids formed on a non-adhesive surface and incubated with 10 µM 5-FU decreased by approximately 1.5-fold, and those incubated with 2 µM 5-FU decreased by 1.1-fold compared to the control (Figure 3B). Smaller spheroids formed using the hanging drop technique and incubated with 10 µM 5-FU decreased by approximately 1.1-fold, while no significant change in diameter was observed after incubation with 2 µM 5-FU (Figure 3B).

When comparing the effects of DOX and 5-FU on smaller spheroids formed using different methods, it was found that, in spheroids formed by the non-adhesive surface method, both DOX concentrations and 10 µM 5-FU inhibited growth 2 to 8 times more strongly than in spheroids formed by the hanging drop method. Meanwhile, the effect of 2 µM 5-FU on spheroid growth did not differ.

Both drugs inhibited the growth of larger (400 µm diameter) A375 spheroids less than that of smaller spheroids (Figure 3). None of the compounds tested statistically significantly inhibited the growth of spheroids formed by the hanging drop technique compared to the control. Also, both 5-FU concentrations had no significant effect on the growth of spheroids formed by both methods (Figure 3D). Spheroids formed on the non-adhesive surface and incubated for eight days with 2 µM and 10 µM DOX had a diameter approximately 1.4-fold smaller than the control (Figure 3C).

When comparing the effects of DOX and 5-FU on the growth of larger spheroids formed using different techniques, it was found that both DOX concentrations inhibited the growth of spheroids formed on the non-adhesive surface by 9 to 22 times more than spheroids formed using the hanging drop technique. Meanwhile, the effect of 5-FU did not differ significantly between spheroids formed by different methods.

The photos show how the size of A375 spheroids changed after eight days of incubation with anticancer compounds compared to the control (Figure 3). Larger and smaller spheroids incubated with DOX grew more slowly, lost their regular spheroid shape, and a layer of adherent cells was observed around the spheroids (Figures 3A and 3C). Larger and smaller spheroids incubated with 10 µM 5-FU remained round in shape (Figures 3B and 3D). After incubation with 2 µM 5-FU, the spheroids visually resembled the control.

The diameter of smaller IGR39 spheroids (200 µm in diameter) formed by both methods and incubated with both DOX concentrations was smaller than that of the control (Figure 4A). Spheroids formed on the non-adhesive surface and incubated with 10 µM DOX had a diameter approximately 1.5-fold smaller, while spheroids formed by the hanging drop technique had a diameter 1.2-fold smaller compared to the control (Figure 4A). After eight days of incubation with 2 µM DOX, smaller spheroids formed on the non-adhesive surface decreased by approximately 1.3-fold, and those formed using the hanging drop technique decreased by 1.2-fold compared to the control (Figure 4A). In summary, both DOX concentrations inhibited the growth of IGR39 spheroids formed by both methods to a similar extent.

**Figure 4.**
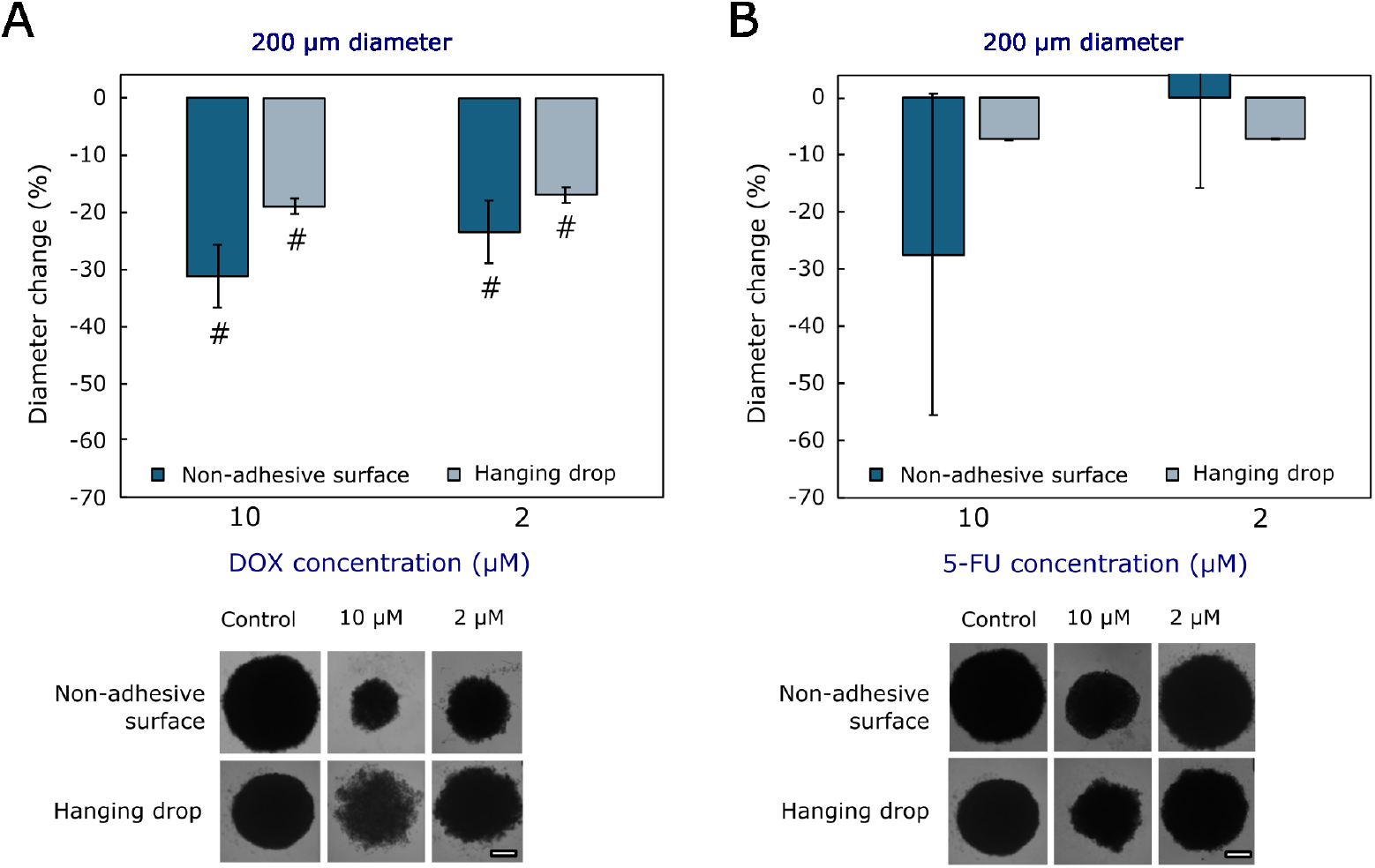
Change in the diameter of 200 µm IGR39 spheroids after 8 days of incubation with DOX (**A**) and 5-FU (**B**); *p < 0.05, statistically significant when comparing spheroids formed by different methods, #p < 0.05, statistically significant when compared to the control. Representative photos of IGR39 spheroids after 8 days of incubation with tested compounds are provided below each graph; the scale bar corresponds to 200 µm.

Following incubation with 10 µM 5-FU, the diameter of smaller spheroids formed on the non-adhesive surface decreased by about 1.4-fold, while incubation with 2 µM 5-FU increased it by about 1.2-fold compared to the control, but no statistically significant differences were found (Figure 4B). A similar effect of 5-FU was observed in spheroids generated by both methods.

Also, it was observed that spheroids formed by both techniques and incubated with both DOX and 10 µM 5-FU concentrations lost their shape, grew more slowly, and a layer of adherent cells was visible around the spheroids (Figure 4). Like the A375 spheroids, the IGR39 spheroids formed by both methods and incubated with 2 µM 5-FU retained their regular shape, and no visual differences were observed compared to the control (Figure 4B).

The effects of DOX and 5-FU on larger IGR39 spheroids (400 µm) could not be assessed, as these spheroids disintegrated within 2-4 days of incubation.

According to previous studies, changes in spheroid diameter do not necessarily indicate the activity of the substances under investigation; cell viability must also be assessed [24,25]. Thus, in addition to morphological changes, we also aimed to compare effects on cell viability in spheroids.

The effect of the cytotoxic drug DOX and the cytostatic drug 5-FU on cell viability was similar to their effects on growth when comparing spheroids generated using different methods. When smaller and larger A375 spheroids with 10 µM DOX were examined, their cell viability was extremely low, ranging from 2 to 2.5% compared to the control (Figures 5A and 5C). In smaller (200 µm diameter) A375 spheroids formed using the hanging drop technique and incubated with 2 µM DOX, cell viability reached about 23% on day 8, while in spheroids of the same size formed on the non-adhesive surface, viability was about 5% (Figure 5A). Meanwhile, the viability of larger (400 µm diameter) spheroids was affected differently by 2 µM DOX, depending on the formation technique: the viability of spheroids formed using the non-adhesive surface method was about 19%, while that of spheroids formed using the hanging drop technique was about 9%. (Figure 5C). These results showed that although the larger A375 spheroids formed by the hanging drop technique were larger, their viability after incubation with 2 µM DOX was lower compared to the spheroids formed on the non-adhesive surface.

**Figure 5.**
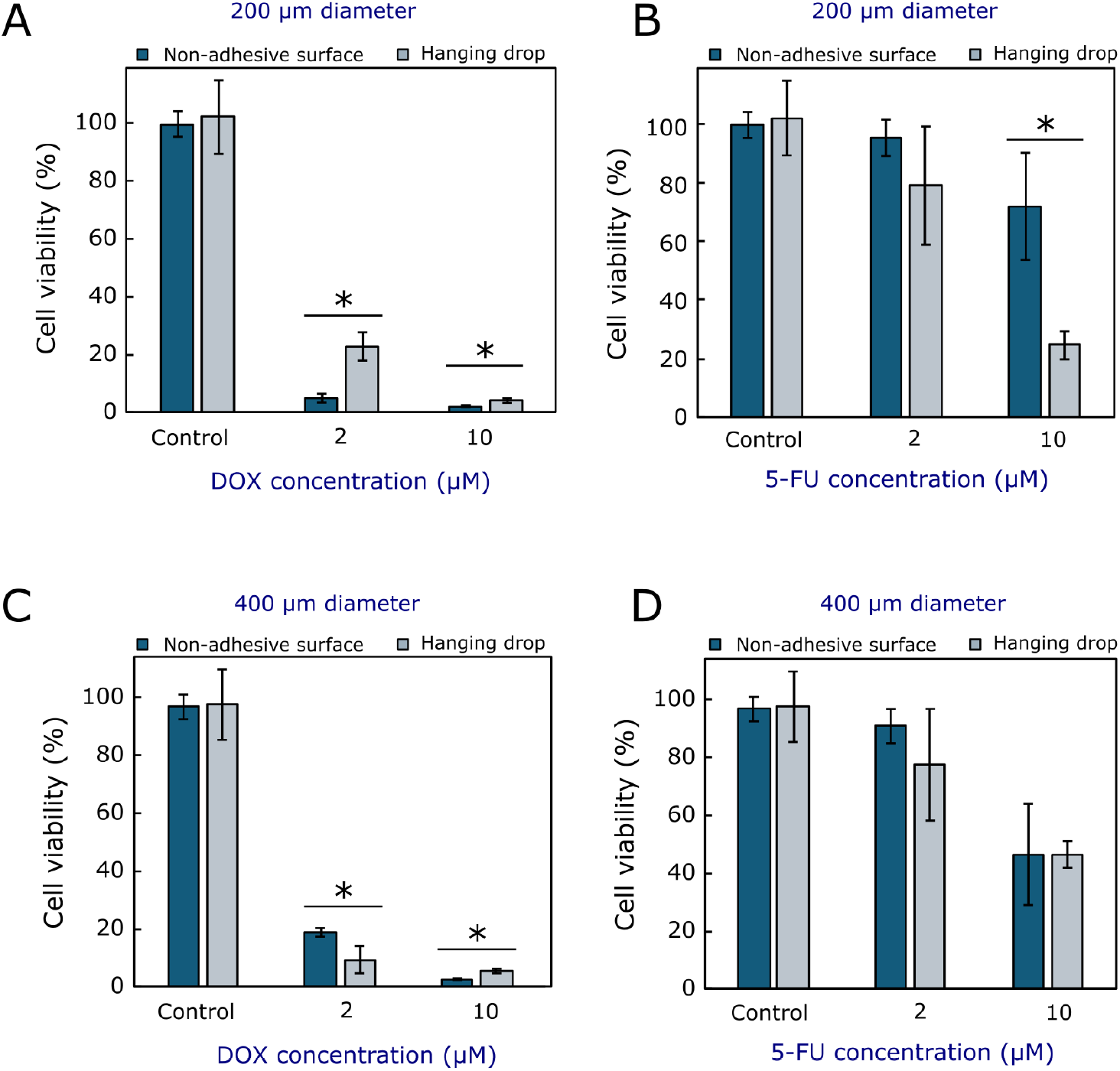
Viability of 200 µm (A, B) and 400 µm (C, D) A375 spheroids after 8 days of incubation with DOX (**A, C**) and 5-FU (**B, D**); *p < 0.05, statistically significant when comparing spheroids formed by different techniques.

Incubation with 2 µM 5-FU did not result in a significant difference in viability between smaller and larger spheroids generated by (Figures 5B and 5D). In smaller A375 spheroids formed on a non-adhesive surface and incubated with 10 µM 5-FU, cell viability was approximately 72%, while in those formed using the hanging drop technique, it was approximately 25% (Figure 5B). In larger spheroids formed by both methods, cell viability was similar (approximately 48%) when incubated with 10 µM 5-FU (Figure 5D).

IGR39 spheroids formed by different methods and incubated with both drug concentrations showed similar cell viability, except when incubated with 2 µM 5-FU (Figure 6). Spheroids formed on the non-adhesive surface and incubated with 2 µM 5-FU retained approximately 95% viability, while those formed by the hanging drop technique retained approximately 64% viability. (Figure 6B). The viability of IGR39 spheroids formed by both techniques was reduced by 10 µM DOX from 3 to 3.7% (Figure 6A) and by 2 µM DOX from 16 to 17% (Figure 6A). After incubation with 10 µM 5-FU, cell viability ranged from 11 to 22% (Figure 6B).

**Figure 6.**
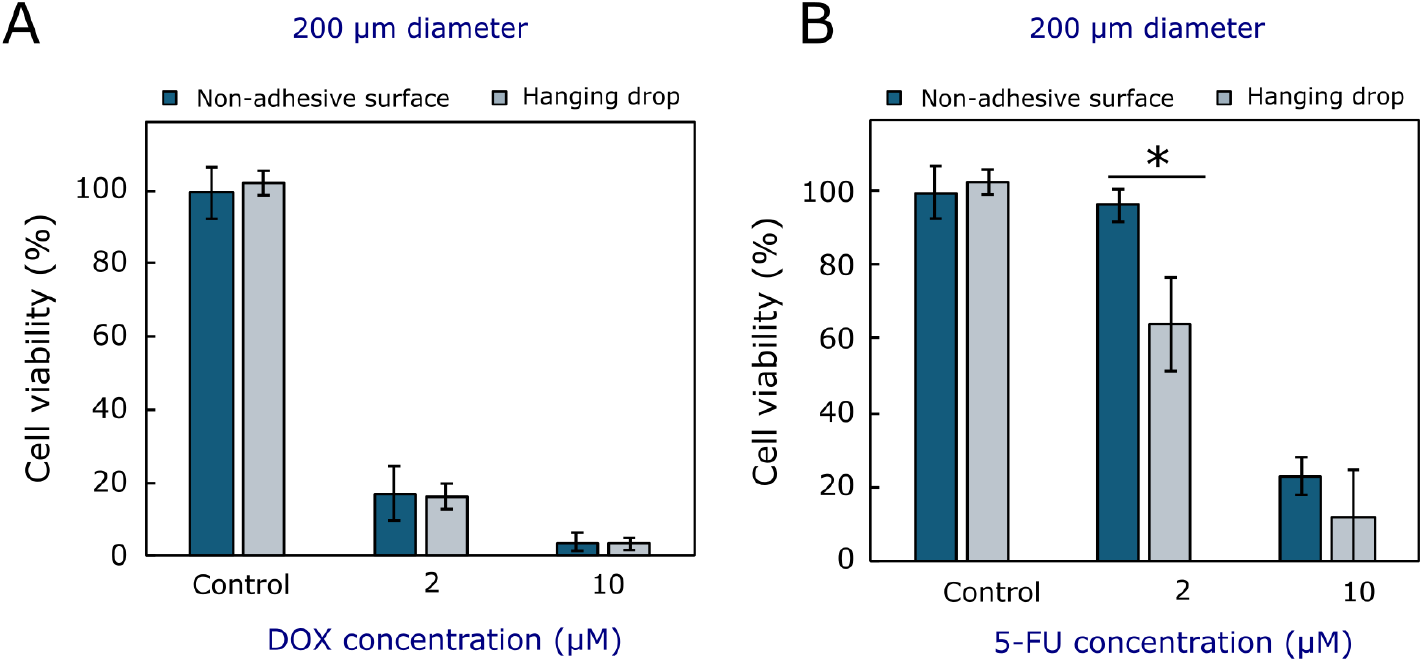
Viability of 200 µm IGR39 spheroids after 8 days of incubation with DOX (**A**) and 5-FU (**B**); *p < 0.05, statistically significant when comparing spheroids formed by different techniques.

Other scientists have also studied and compared the effects of different compounds on spheroid growth and viability, but we were unable to find any published studies comparing the effects on spheroids formed using different techniques, which demonstrates the novelty of our study.

Tayhan investigated how breast cancer spheroids change when incubated with the anticancer drugs DOX and 5-FU and a combination of these drugs. It was found that spheroids incubated with 7 µM DOX or 5 µM 5-FU were approximately 1.1 times smaller than the control after only 24 hours, and the effect of DOX was greater than that of 5-FU, as in our study [26]. Bilgin incubated spheroids formed from different lines of colon cancer, using the non-adhesive surface method, with 2 µM DOX, 200 µM 5-FU, and a combination of these drugs. After 5 days of incubation with 200 µM 5-FU, the diameter of Caco-2 spheroids decreased by 24%, while with 2 µM DOX, the diameter decreased by 21%. The diameter of HT-29 cell spheroids decreased by about 22% after five days of incubation with DOX and 5-FU. Our study also found that after incubation with DOX, the diameter of the spheroids was smaller compared to the control. In the experiment discussed, the researchers found that spheroids incubated with 200 µM 5-FU grew more slowly than spheroids incubated with 2 µM DOX [27].

Valente et al. analysed the effect of DOX on the viability of colon cancer spheroids of the same size (approximately 200 µm and 400 µm) formed on a non-adhesive surface. When these spheroids were incubated with 8 µM DOX, cell viability decreased 2-fold after five days compared to the control [28]. In our study, cell viability was lower than in this study when incubated with higher and lower DOX concentrations.

Mashinchian et al. investigated the effect of 5-FU on breast cancer spheroids formed on a non-adhesive surface. It was found that 50 µM 5-FU reduced viability by approximately 40% after 3 days of incubation. The results of this study indicate that 5-FU has a strong cytotoxic effect on 3D cancer models [29]. In our study, when incubated for longer periods and at lower concentrations of 5-FU, spheroids of a certain size had higher viability compared to those in this study.

Generally, it was found that when smaller and larger A375 spheroids were incubated with 2 and 10 µM DOX, the effect of this drug on spheroid growth and viability depended on the formation method. Spheroids formed on the non-adhesive surface were more sensitive to DOX than spheroids formed using the hanging drop technique. The effect of the cytostatic compound 5-FU on larger A375 spheroids did not depend on the formation method. Smaller spheroids formed using the non-adhesive method became smaller when incubated with 10 µM 5-FU, but were more viable compared to spheroids formed by the hanging drop technique. In IGR39 spheroids, the effect of DOX and 5-FU on their growth did not depend on the formation technique, but 2 µM 5-FU inhibited the viability of spheroids formed on the non-adhesive surface more than spheroids formed by the hanging drop technique.

Summarising all the results of this work, it can be concluded that A375 spheroids formed by the hanging drop and non-adhesive surface methods grew at a similar rate, although at the beginning of the study, spheroids formed by the non-adhesive surface method were larger. Smaller spheroids grew faster than larger ones, regardless of the formation method. IGR39 spheroids formed using the non-adhesive surface method had a smaller diameter than those formed using the hanging drop method, but statistically significant differences were only found during certain incubation periods and with 500 and 1000 cells.

When evaluating the effects of DOX and 5-FU, it was found that A375 spheroids formed on the non-adhesive surface were more sensitive to DOX than spheroids formed using the hanging drop technique. Meanwhile, the effect of 5-FU did not cause a statistically significant change, except for smaller A375 spheroids formed using the non-adhesive surface method. IGR39 spheroids did not show a statistically significant change in size and viability when incubated with DOX and 5-FU, except for spheroids formed using the hanging drop technique, incubated with 2 µM 5-FU, which had lower viability than spheroids formed on the non-adhesive surface.

## 4. Conclusions

The growth dynamics of human melanoma A375 spheroids did not depend on the initial number of cells (from 250 to 2000 cells) or on the method used for their production: formation in a hanging drop or on a non-adhesive surface. When melanoma IGR39 spheroids were formed from 500 and 1000 cells, the spheroids in the hanging drops remained larger than those formed on a non-adhesive surface over a period of ten days. A375 spheroids formed on the non-adhesive surface were more sensitive to DOX than spheroids formed using the hanging drop technique. The effect of 5-FU varied depending on the concentration and size of the spheroids: for larger (400 µm diameter) A375 spheroids, the effect of 5-FU did not depend on the formation technique, while in smaller spheroids (200 µm diameter), 10 µM 5-FU reduced viability more when they were formed using the hanging drop technique. In IGR39 spheroids, the effect of DOX and 5-FU on growth and viability does not depend on the formation method. Based on these results, it can be concluded that the researchers should carefully select the spheroid formation method for their studies, as this may influence the results of the tested compound’s effect on their size and viability.

## Author Contributions

Conceptualisation, V.P.; methodology, V.P; formal analysis, A.Ž.; investigation, A.Ž.; writing—original draft preparation, A.Ž. and V.P.; writing—review and editing, A.Ž. and V.P.; visualisation, A.Ž. and V.P.; supervision, V.P. All authors have read and agreed to the published version of the manuscript.

## Institutional Review Board Statement

This study was conducted in accordance with institutional guidelines and approved by the Bioethical Centre of Lithuanian University of Health Sciences (No. 2024-BEC2-1030; 11 November 2024)

## Informed Consent Statement

Not applicable.

## Data Availability Statement

The original contributions presented in this study are included in the article. Further inquiries can be directed to the corresponding author.

## Abbreviations

2D: Two-dimensional
3D: Three-dimensional
3R: Replacement, reduction, refinement
5-FU: 5-fluorouracil
ATCC: American Type Culture Collection
DMEM
GlutaMAX: Dulbecco’s Modified Eagle GlutaMAX cell culture medium
DMSO: Dimethylsulfoxide
DOX: Doxorubicin
DSMZ: Deutsche Sammlung von Mikroorganismen und Zellkulturen GmbH
ECIS: European Cancer Information System
ER: Estrogen receptors
FBS: Fetal bovine serum
HIF: Hypoxia inducible factor
MC: Methylcellulose
MTT: 3-(4,5-dimethylsthiazol-2-yl)-2,5-diphenyltetrazolium bromide
PBS: Phosphate buffered saline

## References

1. Raghavan, S.; Mehta, P.; Horst, E.N.; Ward, M.R.; Rowley, K.R.; Mehta, G. Comparative Analysis of Tumor Spheroid Generation Techniques for Differential *in Vitro* Drug Toxicity. Oncotarget 2016, 7, 16948–16961, doi:10.18632/oncotarget.7659.

2. Xu, H.; Jiao, D.; Liu, A.; Wu, K. Tumor Organoids: Applications in Cancer Modeling and Potentials in Precision Medicine. J Hematol Oncol 2022, 15, 58, doi:10.1186/s13045-022-01278-4.

3. Gunti, S.; Hoke, A.T.K.; Vu, K.P.; London, N.R. Organoid and Spheroid Tumor Models: Techniques and Applications. Cancers 2021, 13, 874, doi:10.3390/cancers13040874.

4. Zhuang, P.; Chiang, Y.-H.; Fernanda, M.S.; He, M. Using Spheroids as Building Blocks Towards 3D Bioprinting of Tumor Microenvironment. IJB 2024, 7, 444, doi:10.18063/ijb.v7i4.444.

5. Poh, W.T.; Stanslas, J. The New Paradigm in Animal Testing – “3Rs Alternatives.” Regulatory Toxicology and Pharmacology 2024, 153, 105705, doi:10.1016/j.yrtph.2024.105705.

6. Mitrakas, A.G.; Tsolou, A.; Didaskalou, S.; Karkaletsou, L.; Efstathiou, C.; Eftalitsidis, E.; Marmanis, K.; Koffa, M. Applications and Advances of Multicellular Tumor Spheroids: Challenges in Their Development and Analysis. IJMS 2023, 24, 6949, doi:10.3390/ijms24086949.

7. Barbosa, M.A.G.; Xavier, C.P.R.; Pereira, R.F.; Petrikaitė, V.; Vasconcelos, M.H. 3D Cell Culture Models as Recapitulators of the Tumor Microenvironment for the Screening of Anti-Cancer Drugs. Cancers 2021, 14, 190, doi:10.3390/cancers14010190.

8. Daunys, S.; Janonienė, A.; Januškevičienė, I.; Paškevičiūtė, M.; Petrikaitė, V. 3D Tumor Spheroid Models for In Vitro Therapeutic Screening of Nanoparticles. Adv Exp Med Biol 2021, 1295, 243–270, doi:10.1007/978-3-030-58174-9_11.

9. Riffle, S.; Hegde, R.S. Modeling Tumor Cell Adaptations to Hypoxia in Multicellular Tumor Spheroids. J Exp Clin Cancer Res 2017, 36, 102, doi:10.1186/s13046-017-0570-9.

10. Vakhshiteh, F.; Bagheri, Z.; Soleimani, M.; Ahvaraki, A.; Pournemat, P.; Alavi, S.E.; Madjd, Z. Heterotypic Tumor Spheroids: A Platform for Nanomedicine Evaluation. J Nanobiotechnol 2023, 21, 249, doi:10.1186/s12951-023-02021-y.

11. Ostrowski, S.M.; Fisher, D.E. Biology of Melanoma. Hematology/Oncology Clinics of North America 2021, 35, 29–56, doi:10.1016/j.hoc.2020.08.010.

12. Nurla, L.A.; Forsea, A.-M. Melanoma Epidemiology in Europe: What Is New? Ital J Dermatol Venereol 2024, 159, doi:10.23736/S2784-8671.24.07811-3.

13. Živković, Z.; Opačak-Bernardi, T. An Overview on Spheroid and Organoid Models in Applied Studies. Sci 2025, 7, 27, doi:10.3390/sci7010027.

14. Tevlek, A.; Kecili, S.; Ozcelik, O.S.; Kulah, H.; Tekin, H.C. Spheroid Engineering in Microfluidic Devices. ACS Omega 2023, 8, 3630–3649, doi:10.1021/acsomega.2c06052.

15. Lv, J.; Du, X.; Wang, M.; Su, J.; Wei, Y.; Xu, C. Construction of Tumor Organoids and Their Application to Cancer Research and Therapy. Theranostics 2024, 14, 1101–1125, doi:10.7150/thno.91362.

16. Loo, E.; Khalili, P.; Beuhler, K.; Siddiqi, I.; Vasef, M.A. BRAF V600E Mutation Across Multiple Tumor Types: Correlation Between DNA-Based Sequencing and Mutation-Specific Immunohistochemistry. Applied Immunohistochemistry & Molecular Morphology 2018, 26, 709–713, doi:10.1097/PAI.0000000000000516.

17. Marzagalli, M.; Casati, L.; Moretti, R.M.; Montagnani Marelli, M.; Limonta, P. Estrogen Receptor β Agonists Differentially Affect the Growth of Human Melanoma Cell Lines. PLoS ONE 2015, 10, e0134396, doi:10.1371/journal.pone.0134396.

18. Pefani-Antimisiari, K.; Athanasopoulos, D.K.; Marazioti, A.; Sklias, K.; Rodi, M.; De Lastic, A.-L.; Mouzaki, A.; Svarnas, P.; Antimisiaris, S.G. Synergistic Effect of Cold Atmospheric Pressure Plasma and Free or Liposomal Doxorubicin on Melanoma Cells. Sci Rep 2021, 11, 14788, doi:10.1038/s41598-021-94130-7.

19. Koleini, N.; Kardami, E. Autophagy and Mitophagy in the Context of Doxorubicin-Induced Cardiotoxicity. Oncotarget 2017, 8, 46663–46680, doi:10.18632/oncotarget.16944.

20. Salvador, D.; Bastos, V.; Oliveira, H. Hyperthermia Enhances Doxorubicin Therapeutic Efficacy against A375 and MNT-1 Melanoma Cells. IJMS 2021, 23, 35, doi:10.3390/ijms23010035.

21. Milczarek, M.; Pogorzelska, A.; Wiktorska, K. Synergistic Interaction between 5-FU and an Analog of Sulforaphane—2-Oxohexyl Isothiocyanate—In an In Vitro Colon Cancer Model. Molecules 2021, 26, 3019, doi:10.3390/molecules26103019.

22. Tian, J.; Zhang, D.; Kurbatov, V.; Wang, Q.; Wang, Y.; Fang, D.; Wu, L.; Bosenberg, M.; Muzumdar, M.D.; Khan, S.; et al. 5-Fluorouracil Efficacy Requires Anti-tumor Immunity Triggered by Cancer-cell-intrinsic STING. The EMBO Journal 2021, 40, e106065, doi:10.15252/embj.2020106065.

23. Jeong, Y.; Tin, A.; Irudayaraj, J. Flipped Well-Plate Hanging-Drop Technique for Growing Three-Dimensional Tumors. Front. Bioeng. Biotechnol. 2022, 10, 898699, doi:10.3389/fbioe.2022.898699.

24. Pranaitytė, G.; Grybaitė, B.; Endriulaityte, U.; Mickevičius, V.; Petrikaitė, V. Exploration of 1-(2,4-Difluorophenyl)-5-Oxopyrrolidine-3-Carboxylic Acid Derivatives Effect on Triple-Negative Breast, Prostate Cancer and Melanoma Cell 2D and 3D Cultures. Sci Rep 2025, 15, 17590, doi:10.1038/s41598-025-02106-8.

25. Kulesza, J.; Pawłowska, M.; Augustin, E. The Influence of Antitumor Unsymmetrical Bisacridines on 3D Cancer Spheroids Growth and Viability. Molecules 2021, 26, 6262, doi:10.3390/molecules26206262.

26. Erden Tayhan, S. A Study with Cancer Stem Cells and Three-Dimensional Tumoroids: Investigation of the Combined Effects of 5-Fluorouracil and Doxorubicin in Breast Cancer. Med Oncol 2024, 41, 185, doi:10.1007/s12032-024-02423-4.

27. Bilgin, S. Apoptotic Effect of 5-Fluorouracil-Doxorubicin Combination on Colorectal Cancer Cell Monolayers and Spheroids. Mol Biol Rep 2024, 51, 603, doi:10.1007/s11033-024-09562-x.

28. Valente, R.; Cordeiro, S.; Luz, A.; Melo, M.C.; Rodrigues, C.R.; Baptista, P.V.; Fernandes, A.R. Doxorubicin-Sensitive and -Resistant Colorectal Cancer Spheroid Models: Assessing Tumor Microenvironment Features for Therapeutic Modulation. Front. Cell Dev. Biol. 2023, 11, 1310397, doi:10.3389/fcell.2023.1310397.

29. Mashinchian, M.; Vandghanooni, S.; Karamibonari, A.R.; Eskandani, M. β-Hydroxybutyrate Promotes Chemoresistance and Proliferation in Breast Cancer Cells. Biochemistry and Biophysics Reports 2025, 44, 102217, doi:10.1016/j.bbrep.2025.102217.

